# Single cell RNA-seq and ATAC-seq indicate critical roles of Isl1 and Nkx2-5 for cardiac progenitor cell transition states and lineage settlement

**DOI:** 10.1101/210930

**Authors:** Guangshuai Jia, Jens Preussner, Stefan Guenther, Xuejun Yuan, Michail Yekelchyk, Carsten Kuenne, Mario Looso, Yonggang Zhou, Thomas Braun

## Abstract

Formation and segregation of cell lineages building the vertebrate heart have been studied extensively by genetic cell tracing techniques and by analysis of single marker gene expression but the underlying gene regulatory networks driving cell fate transitions during early cardiogenesis are only partially understood. Here, we comprehensively characterized mouse cardiac progenitor cells (CPC) marked by Nkx2-5 and Isl1 expression from E7.5 to E9.5 using single-cell RNA sequencing. By leveraging on cell-to-cell heterogeneity, we identified different previously unknown cardiac sub-populations. Reconstruction of the developmental trajectory revealed that Isl1^+^ CPC represent a transitional cell population maintaining a prolonged multipotent state, whereas extended expression of Nkx2-5 commits CPC to a unidirectional cardiomyocyte fate. Furthermore, we show that CPC fate transitions are associated with distinct open chromatin states, which critically depend on Isl1 and Nkx2-5. Our data provide a model of transcriptional and epigenetic regulations during cardiac progenitor cell fate decisions at single-cell resolution.

---

Cell fate mapping experiments demonstrated that cardiac progenitor cells (CPC) in the mouse form from Mesp1^+^ cells that leave the primitive streak during gastrulation at E6.5 (reviewed by [1]). At E7.5, CPC express the homeobox genes Nkx2-5, Isl1, or a combination of both and exhibit a multilineage potential enabling them to generate cardiomyocytes, smooth muscle cells, endothelial cells and pericytes [2, 3]. During early developmental stages CPC are present in two distinct anatomical locations, the first (FHF) and the second heart field (SHF) [4–6]. Unlike FHF cells, SHF cells show delayed differentiation into myocardial cells and represent a reservoir of multipotent CPC during cardiogenesis (reviewed by [1]). Isl1 is expressed in cardiac progenitor cells (CPC) of the SHF, although a broader, transient expression has been noted in the anterior intra-embryonic coelomic walls and proximal head mesenchyme encompassing both the FHF and the SHF [7, 8]. However, efficient nGFP labeling of CPC is only achieved in the SHF, making the Isl1^nGFP/+^ knock-in reporter mouse line a reliable source for isolation of SHF cells [9, 10]. In contrast, Nkx2-5 expression marks cells of both the FHF and SHF including the cardiac crescent and the pharyngeal mesoderm [1, 7, 11]. Although transient co-expression of Isl1 and Nkx2-5 has been observed, several lines of evidence indicate that Isl1 and Nkx2-5 suppress each other thereby allowing expansion of Isl1^+^ CPC and differentiation into Nkx2-5^+^ cardiomyocytes [7, 10].

Differentiated cells (e.g. cardiomyocytes) are assumed to acquire their identity in a successive step-wise manner from multipotent cells (e.g. CPC) but the different intermediate states allowing transition from multipotent precursor cells to differentiated descendants still await further characterization. Global analysis of transcriptional changes does not provide the resolution to precisely identify such specific cellular transition states. However, recent advances in single-cell RNA sequencing (scRNA-seq) permit characterization of transcriptomes at the single cell level at multiple time points, thereby enabling detailed assessment of developmental trajectories of precursor cells [12].

Analysis of large groups of closely related transcriptomes pose specific challenges for computational methods, which can be met by projection of data points from the high-dimensional gene expression space into a low-dimensional latent space [13, 14][15–18]. Inclusion of landmark genes [19] and microdissection of relevant structures of murine hearts have further refined this approach [20, 21]. Furthermore, pseudotemporal ordering [22] has been adapted for scRNA-seq data [23, 24] to reveal cell fate transitions throughout developmental processes [25]. Transcriptional changes are either preceded, followed, or accompanied by changes in chromatin organization [26]. ATAC-seq (an assay for transposase-accessible chromatin using sequencing) provides a robust means to identify chromatin closure and opening at enhancers and promoters in a limited number of cells [27] but has not been applied yet to characterize chromatin accessibility and putative regulatory elements driving cardiogenesis.

Here, we used scRNA-seq to transcriptionally profile FACS-purified Nkx2-5^+^ and Isl1^+^ cells from E7.5, E8.5 and E9.5 mouse embryos. We decided to focus on native embryonic cells and not on ESC derivatives, since *in vitro* results have to be viewed with caution despite some advantages of ESC-based approaches [28, 29]. By taking advantage of unsupervised bioinformatics analysis, we reconstructed the developmental trajectories of Nkx2-5^+^ and Isl1^+^ cells. We found that Isl1^+^ CPC behaves as a transitory cell population, which become developmentally arrested after inactivation of Isl1. We show that forced expression of Nkx2-5 primes the cardiomyocyte fate and used ATAC-seq to characterize accompanying changes in the chromatin landscape. Our study provides a rich source for future studies aiming to dissect the functional role of newly identified genes in cardiac development and congenital heart disease.

## RESULTS

### Single cell transcriptomics reveals heterogeneity of cardiac progenitor cells

To unravel the molecular composition of either Isl1^+^ or Nkx2-5^+^ CPC, we isolated GFP^+^ cells by FACS from Nkx2-5-emGFP transgenic and Isl1^nGFP/+^ knock-in embryos (Fig. 1a) at E7.5, E8.5 and E9.5 and performed single-cell RNA sequencing using the Fludigm C1 workstation (Fig. 1b). At E8.5 and E9.5 we used dissected hearts instead of the whole embryo, to avoid contamination of non-cardiogenic cells that might be marked by Isl1 or Nkx2-5 expression. After removal of low quality cells (Supplementary Fig. 1a-g), we obtained 167 Nkx2-5^+^ and 254 Isl1^+^ cell transcriptomes, which cover most stages of early heart development (Fig. 1b).

**Figure 1.**
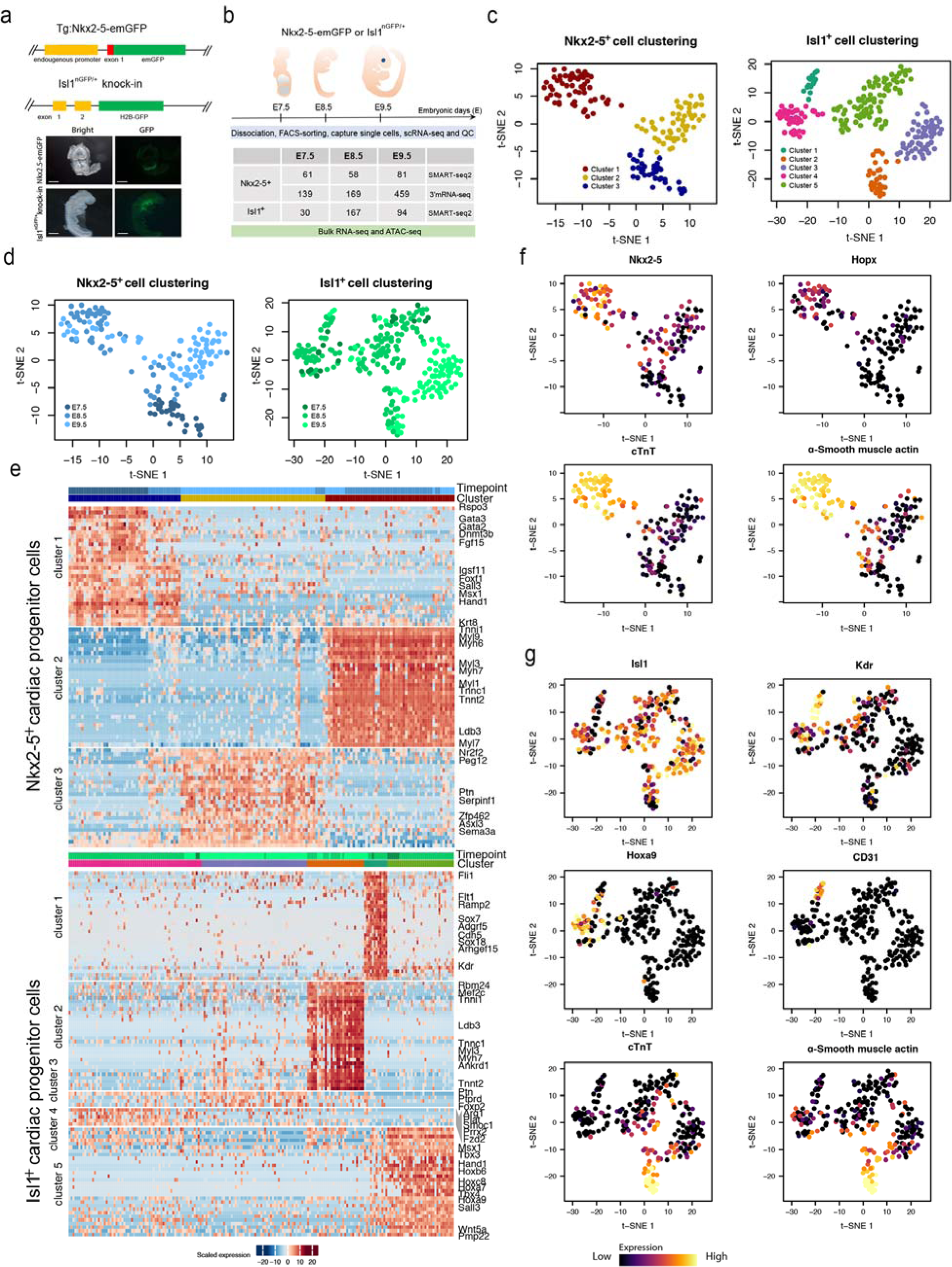
Single-cell RNA-seq of CPC and identification of heterogeneous subpopulations and maker genes. (**a**) Schematic representation of the Nkx2-5-emGFP transgenic reporter and Isl1^nGFP/+^ allele (top). Expression of Nkx2-5-emGFP and Isl1-nGFP at E8.5 in mouse embryonic hearts. (bottom). (**b**) Sampling time points for scRNA-seq, bulk RNA-seq, ATAC-seq. The table shows numbers of cells used for scRNA-seq. QC: quality control. (**c**) t-SNE visualization of individual Nkx2-5^+^ and Isl1^+^ CPC to identify subpopulations. Colors denote corresponding clusters, and (**d**) development stages. (**e**) Hierarchical clustering of expression heatmaps showing differentially expressed marker genes (AUROC > 0.8, FDR < 0.01; and lower bound of LogFC > 2 or higher bound of LogFC < −2, FDR < 0.01) across different clusters in Nkx2-5 (top) and Isl1 (bottom). (**f**) Expression of selected individual genes in Nkx2-5^+^ and Isl1^+^ (**g**) cells. The color represents expression levels of cells that are shown in the t-SNE plots in (**c**). (Scale bars: 300 μm).

We first asked whether Nkx2-5^+^ and Isl1^+^ CPC sampled at successive developmental time-points represent homogenous populations or are composed of distinct subpopulations. Therefore, we analyzed the coefficient of variation and dropout rates of defined heterogeneous genes as input for a neuronal network-based dimension reduction strategy (self-organizing map, SOM) [30], (Supplementary Fig. 2a & b). The resulting SOM were projected into 2D for visualization by t-distributed stochastic neighbor embedding (t-SNE). This strategy allowed us to identify three major subpopulations of Nkx2-5^+^ and five subpopulations of Isl1^+^ cells (Fig. 1c). The Nkx2-5^+^ cluster 3 mainly comprised E7.5 cells, whereas cluster 1 contained cells from E8.5 and E9.5 implying an intermediate cell state. Cluster 2 predominantly contained cells from E9.5 (Fig. 1d). Stage-dependent clustering was less evident for the five Isl1^+^ subpopulations, which might indicate that the specific cellular phenotypes of Isl1^+^ subpopulations are maintained for longer time periods even in a changing developmental environment (Fig. 1d).

To identify cluster-specific, differentially expressed genes, we used MAST [31] and a gene ranking approach implemented in SC3 [32]. The top 269 and 216 genes that were differentially expressed in the Nkx2-5^+^ and the Isl1^+^ lineage, respectively, included several established cardiac regulatory genes such as *Hand1, Tbx3/4/5, Gata2/3, Smarcd3, Rbm24, Wnt5a, Bmp4, Notch1* and *Fgf3/15* (Fig. 1e; Supplementary Table 1, 2) [20, 21, 33–35]. Importantly, we detected numerous differentially expressed genes that so far had not been linked to cardiogenesis probably due to restricted expression in a small number of cells, which rendered them invisible to conventional bulk transcriptome analysis (Fig. 1e; Supplementary Table 1, 2). For example, Isl1^+^ cluster 5 expressed the cardiac transcription factors *Tbx3/4* and *Wnt5a* as well as several posterior *Hox* genes including *Hoxa7/9/10, Hoxb6, Hoxc8* and *Hoxd8* (Fig. 1e; Supplementary Fig. 3a), which might contribute to cardiac patterning in the second heart field between E7.5 and E8.5. In contrast to posterior *Hox* genes, which were specifically expressed in Isl1^+^ cluster 5, the anterior Hox genes *Hoxa1, Hoxa3* and *Hoxb1* were active in individual cells scattered across different CPC clusters (Supplementary Fig. 3b). In addition, we newly identified several transcription factors such as *Sox7/18, Sall3, Ldb3, Zbtb20, Zfp462/512b/711, Klf14*; G-proteins including *Arhgap1, Adgrf5, Arhgef15*; and the *de novo* DNA methyltransferase *Dnmt3b* in Nkx2-5^+^ or Isl1^+^ clusters (Fig. 1e; Supplementary Table 1, 2).

Next, we assigned identities to each cluster based on the expression of key marker genes (Fig. 1f, g). Consistent with the gene ontology analysis of differentially expressed genes within each cluster, Nkx2-5^+^ cluster 3 and Isl1^+^ cluster 2, characterized by *cTnt* and α*-smooth muscle actin* expression appear to represent a myogenic fate whereas Isl1+ cluster 1, expressing *Cd31*, is characterized by endothelial cell features (Fig. 1f, g; Supplementary Fig. 4). Interestingly, expression of *Nkx2-5* and *Isl1* varied among subpopulations within each lineage: (i) Nkx2-5 shows more pronounced expression in late stages (clusters 3 and 1, E8.5 and E9.5) (Fig. 1f, Fig. 2e); (ii) Isl1 expression decreases in the cells expressing differentiation markers (clusters 2 and 1) (Fig. 1g, Fig. 2f). This pattern suggests that Nkx2-5 is associated with myogenic differentiation while Isl1 is linked to the maintenance of progenitor cell multipotency, which is consistent with previous models [10, 36, 37].

**Figure 2.**
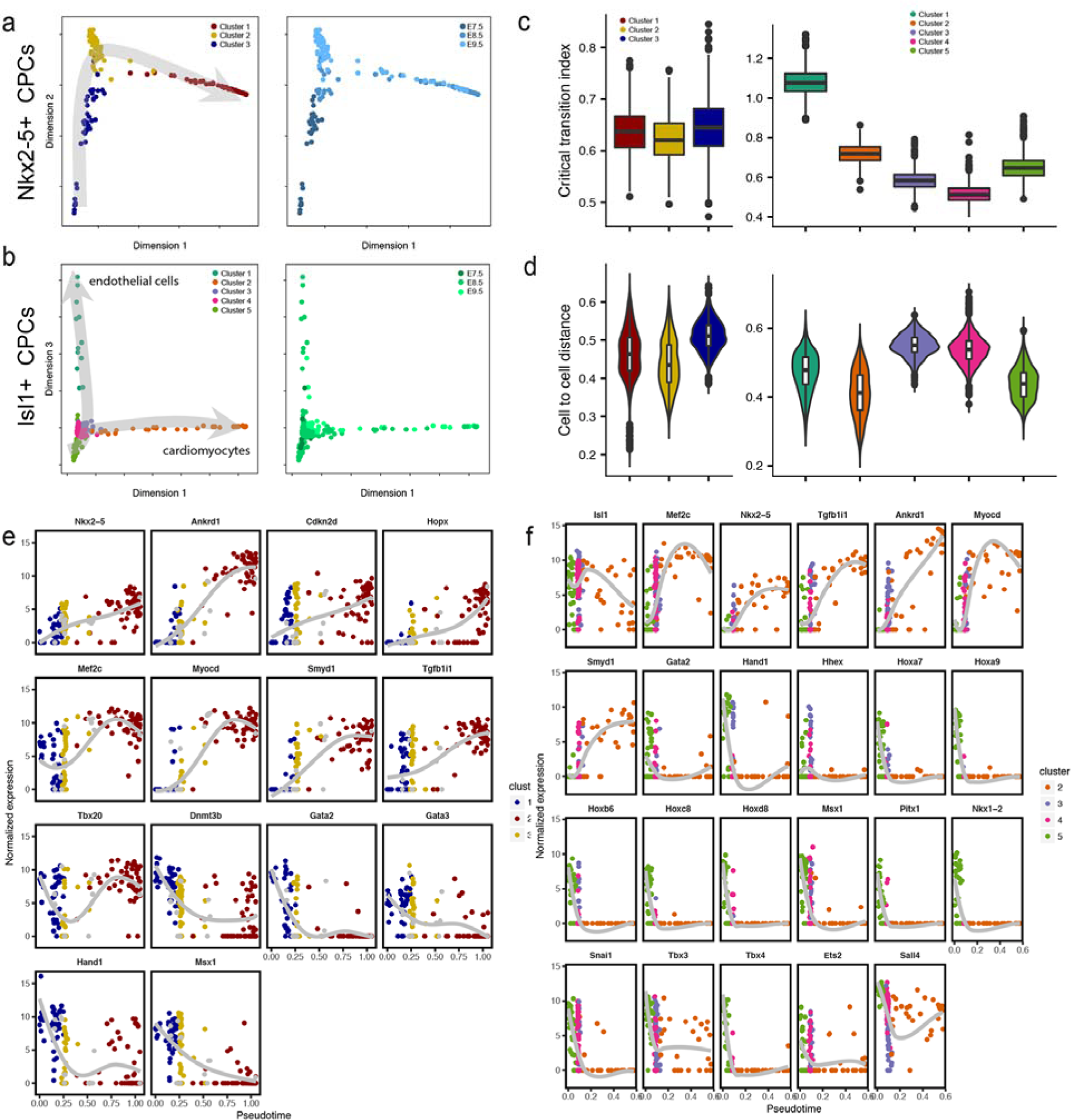
Reconstruction of developmental trajectories of cardiac progenitor cells and computation of transition states. (**a**) t-SNE plots showing the diffusion pseudotimes of Nkx2-5^+^ and (**b**) Isl1^+^ CPC. Clusters and development stages of individual cells are colored coded as indicated. (**c**) Boxplots representing distribution of I_C_(C) values from all marker genes of each cluster of Nkx2-5^+^ (left) and Isl1^+^ (right) cells. P-value for comparison between clusters (Kolmogorov–Smirnov test and Wilcoxon rank sum test). (**d**) Boxplots showing pairwise cell-to-cell distances across each cluster of Nkx2-5^+^ (left) and Isl1^+^ (right) cells. (**e, f**) Expression levels of different transcription factors and key marker genes along the pseudotime axis in Nkx2-5^+^ (**e**) and Isl1^+^ (**f**) cells.

To test the robustness of our approach and to analyze whether we have sequenced sufficient numbers of cells to unveil the entire heterogeneity of CPC, we generated single-cell transcriptomes of additional 663 Nkx2-5^+^ CPC using WaferGen iCell8 system (Fig. 1b). To correct for the resulting batch effects, which in part are due to differences in the sequencing depth of libraries generated with C1 and iCell8 systems (Supplementary Fig. 1a-e), we applied the MNN method [38]. After successful merging and aligning data from both approaches (Supplementary Fig. 5a, b), we performed the same analysis as described above. We identified 3 clusters of Nxk2-5^+^ CPC, essentially mirroring the C1 data (Supplementary Fig. 5c, d). The consistent recapitulation of subpopulations when using significantly more cells (663 vs 167 cells) suggests that even comparatively low numbers are sufficient to unravel the heterogeneity among Nxk2-5^+^ CPC. Furthermore, we compared genes that were found to be differentially expressed among subpopulations using the C1 platform with the merged data set. We detected a similar distribution of marker genes in cluster 1 and 3 but did not fully reproduce the marker gene pattern for cluster 2 (Supplementary Fig. 5e). We concluded that sequencing depth rather than cell numbers is the main limiting factor for discovery of novel genes in cardiac progenitor cells. Thus, we hereafter focused our analysis on the C1 data, which provided substantially deeper sequence coverage (Supplementary Fig. 1a-e).

### Reconstruction of development trajectories reveals differential developmental potential of Isl1^+^ and Nkx2-5^+^ CPC

scRNA-seq data allow ordering of cells by “pseudotime” based on cell-to-cell transcriptome similarity for calculation of developmental trajectories. We took advantage of diffusion maps to arrange cardiac progenitor cells according to their developmental pseudotime [24]. We mapped cells collected at successive developmental stages along the pseudotime for reconstruction of the developmental trajectories of Nkx2-5^+^ and Isl1^+^ CPC (Fig. 2a, b). Interestingly, cells collected at the same embryonic stages aligned to broad pseudotime points, which suggested that CPC are not synchronized at different embryonic stages but follow individual developmental traits. Next, we aligned the different Nkx2-5^+^ and Isl1^+^ cell clusters to developmental trajectories. Nkx2-5^+^ CPC showed one continuous trajectory suggesting a unipotent differentiation capacity (Fig. 2a). In contrast, the trajectory of Isl1^+^ CPC bifurcated into two distinct orientations (endothelial cells and cardiomyocytes), suggesting the existence of a transition state, which separates multipotency of Isl1^+^ CPC from acquisition of distinct cellular identities (Fig. 2b).

Cells undergoing a critical fate decision (such as lineage bifurcation) have been postulated to pass a transition state [25], which corresponds to a switch between different attractor states [39]. To delineate such transition states, we calculated the critical transition index of Nkx2-5^+^ and Isl1^+^ cell clusters [abbreviated as *I_c_(c)*] [40]. The *I_c_(c)* values of Nkx2-5^+^ clusters showed similar numerical ranges essentially excluding existence of a transition state in the Nkx2-5^+^ cell population. Instead, cells from later stages (cluster 1) showed decrease of *I_c_(c)* indicating stable settlement into an attractor state (Fig. 2c, left) with cardiomyocyte-like expression characteristics (Fig. 1e). In contrast, computation of *I_C_(c)* values of Isl1^+^ clusters revealed decreased values for cells at the bifurcation point (cluster 3 and 4), indicating appearance of a transition state. Cells that overcame this point (clusters 1 and 2) exhibited more coordinated changes of gene expression changes (Fig. 2c, right). Since the critical transition index only hints to the presence of a transition state but does not reveal its stability, we calculated pairwise cell-to-cell distances [41] (Supplementary Fig. 6a & b). High cell-to-cell distances in seemingly homogeneous populations indicate transcriptional noise, which occurs when several gene regulatory networks become active opening new opportunities for cellular decisions. As expected, cell-to-cell distances of Nkx2-5^+^ CPC did not change dramatically while cell-to-cell distances of Isl1^+^ CPC in cluster 3 and 4 increased substantially (Fig. 2d). Taken together our results show that Nkx2-5^+^ CPC follow a straight path towards their cardiomyocyte fate whereas Isl1^+^ CPC need to overcome a transition state with elevated noise levels.

To identify genes that are potentially required to determine and/or maintain correspondent cell states, we generated a list of 108 genes for Nkx2-5^+^ cells and 130 genes for Isl1^+^ cells positively correlated with progression of pseudotime across clusters (Spearman rank correlation coefficient > 0.5; Supplementary Fig. 7a, b; Supplementary Table 3) focusing on transcription factors (TFs) and their chromatin-modifying partners (Fig. 2e, f). Furthermore, we assigned genes expressed highly at early stages of the developmental trajectory to a “*priming*” category. Genes expressed at fate-restricted stages but not in multipotent progenitor cells were placed in a “*de novo*” category (Supplementary Fig. 7c, d). We noted that expression of *Dnmt3b, Gata2/3, Hand1* and *Msx1* declined along Nkx2-5^+^ developmental trajectories, qualifying them as “*priming*” genes. In contrast, we noted increased expression of *Nkx2-5, Ankrd1, Cdkn2d, Hopx, Mef2c, Myocd, Smyd1, Tgfb1i1 and Tbx20* during Nkx2-5^+^ CPC differentiation qualifying them as “*de novo*” genes (Fig. 2e). The “p*riming”* genes for Isl1^+^ CPC included transcription factors of the *Hox*-family (*Hoxa7, Hoxa9, Hoxb6, Hoxc8* and *Hoxd8*), *Gata2, Hand1,* and *Tbx3/4* as well as *Hhex, Msx1, Pitx1, Nkx1-2, Ets2, Sall4 and Snai1*. *Mef2c, Nkx2-5, Tgfb1i1, Ankrd1, Myocd* and *Smyd1* were expressed at fate-restricted stages and hence represent “*de novo*” genes (Fig. 2f). The distinct expression pattern of “*priming”* and “*de novo”* TFs in Nkx2-5^+^ compared to Isl1^+^ cells indicates that the fate of progenitor cells is governed by different gene regulatory networks compared to fate-restricted cells.

### Isl1 is indispensable for cardiac progenitor cell fate bifurcation

The loss of Isl1 results in absence of outflow tract and right ventricle and early embryonic lethality [42], which prevents dissection of Isl1-dependent molecular processes in the SHF. To address the role of Isl1 in cell fate determination, we inactivated the *Isl1* gene by generating Isl1^nGFP/nGFP^ embryos and isolated Isl1-GFP^+^ cells by FACS for scRNA-seq analysis at E9.5 (Fig. 3a). Projection of Isl1-KO single cells on the trajectory of the developing SHF revealed that Isl1-KO cells are stalled/trapped in the previously identified stable attractor state (Fig. 3b). Analysis of G1/S and G2/M cell cycle markers in single cells (Supplementary Fig. 8a-c; Methods) suggested a reduction of cycling Isl1-knockout cells compared to wild type cells, although the results did not reach statistical significance (χ^2^ test: *p*= 0.062) (Fig. 3c). We concluded that proliferation defects might contribute to the attractor state of Isl1-knockout cells but that additional biological processes probably play more important roles. To identify such processes, we performed gene ontology analysis of deregulated genes in Isl1-knockout cells. Interestingly, cell differentiation genes were strongly affected by the inactivation of Isl1 (Fig. 3d). Moreover, the GO terms “endothelial cell migration” and “muscle organ development” were enriched in WT compared to Isl1-knockout cells (Fig. 3d), which is consistent with the compromised differentiation of Isl1-knockout cells to endothelial cells or cardiomyocytes.

**Figure 3.**
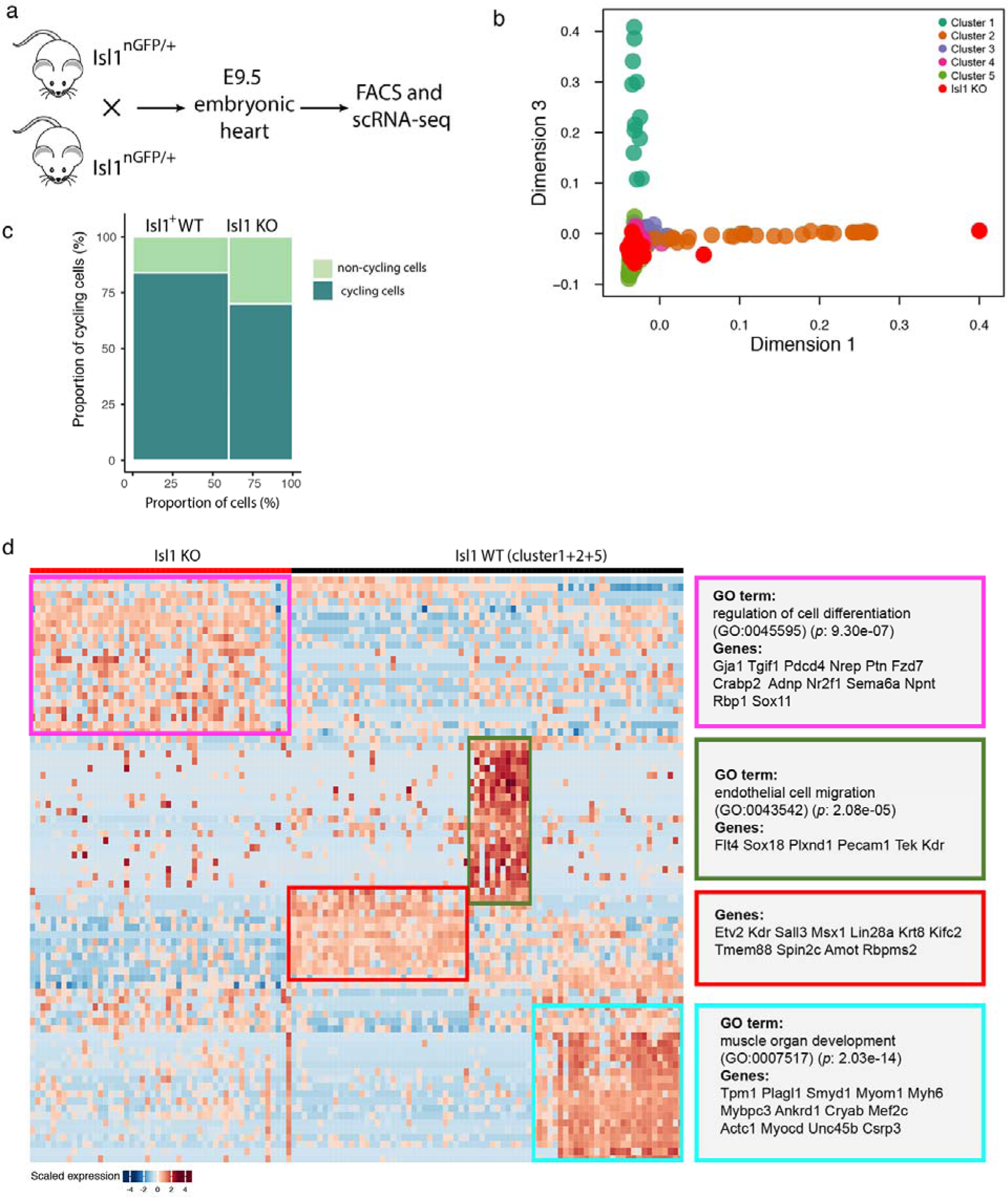
scRNA-seq of Isl1 knockout CPC. (**a**) Schematic illustration depicting generation of Isl1 embryos and scRNA-seq. (**b**) Predicted diffusion pseudotime of Isl1 knockout CPC projected on t-SNE plots of Isl1^+^ cells. (**c**) Ratios of cycling and non-cycling Isl1 knockout and Isl1^+^ wild type CPC. (**d**) Heatmap showing expression of deregulated genes in Isl1^+^ cells at E8.5 and E9.5 (cluster 1, 2 and 5) isolated from Isl1 knockout and control embryos.

### Nkx2-5 establishes a unidirectional fate for CPC towards cardiomyocytes

Our pseudotime-based analysis of developmental trajectories revealed one continuous trajectory of Nkx2-5^+^ CPC suggesting that Nkx2-5^+^ cells are exclusively committed to become cardiomyocytes. Such a conclusion seems incompatible with previous lineage tracing studies, which suggested that Nkx2-5^+^ cells contribute to other cell types such as smooth muscle cells [43]. We reasoned that Nkx2-5 expression is essential to maintain the ability of multipotent progenitor cells to differentiate into cardiomyocytes but that Nkx2-5 expression is quickly terminated in cells, which acquire a stable smooth muscle cell fate thereby escaping Nkx2-5-emGFP based FACS-sorting. scRNA-Seq analysis indicated that Nkx2-5^+^ cells at E8.5 express cardiac markers such as *cTNT* but also *α-SMA* as well as several other smooth muscle markers such as *Caldesmon*, *Tagln* and *Cnn1* (Fig. 1f, g; Supplementary Fig. 9). The co-expression of cardiomyocyte and smooth muscle cell markers might suggest the ability of Nkx2-5^+^ cells to differentiate into cardiomyocytes and smooth muscle cells but might alternatively reflect the well-known expression of smooth muscle genes in immature cardiomyocytes [44]. Cardiac priming at E8.5 due to continued expression of Nkx2-5 might overcome smooth muscle identity and induce a stable cardiomyocyte fate.

To directly test this hypothesis, we first re-analyzed published scRNA-seq data of Nkx2-5 null embryonic hearts at E9.5 [20] and found significantly increased numbers of smooth muscle cells raising from 14.5% (138/949 cells) in wildtype to 31.2% (39/125 cells) in Nkx2-5 mutant hearts (χ^2^ test: *p* < 2.37e-6) (Fig. 4a). In a second approach, we specifically expressed Nkx2-5 and EGFP (separated by an IRES) in the Isl1^+^ lineage using Isl1-Cre to initiate transcription from the Rosa26 locus (hereafter named Isl1^+^/Nkx2-5OE) [10]. Isolation of GFP^+^ cells by FACS from E12.5 embryonic hearts and scRNA-seq (Fig. 4b) revealed that Isl1^+^/Nkx2-5OE cells align to the Nkx2-5^+^ trajectories and the cardiomyocyte-like branch of the Isl1^+^ trajectory (Fig. 4c, d). Importantly, Isl1^+^/Nkx2-5OE cells did not contain any endothelial cell- or smooth muscle cell-like populations, although by E12.5 Isl1^+^ cells have given rise to multiple endothelial cells in wildtype conditions (Fig. 4d). Taken together our results indicate that Nkx2-5 is required and sufficient to resolve the multipotent differentiation capacity of CPC.

**Figure 4.**
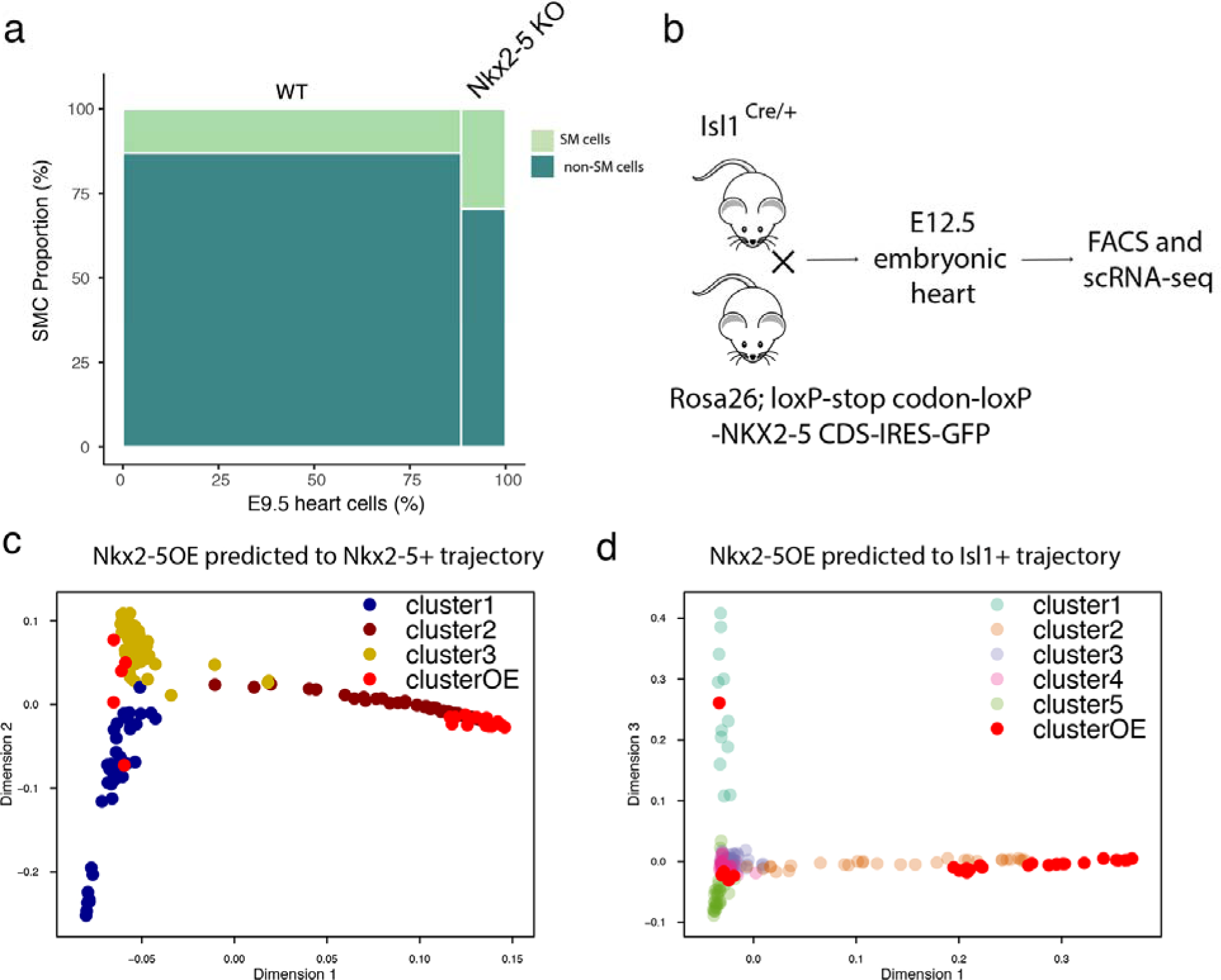
scRNA-seq of Isl1^+^/Nkx2-5OE cells. (**a**) Re-analysis of published data showing the ratio of smooth muscle cells in embryonic hearts of wild type and Nkx2-5 knockout embryos at E9.5. The smooth muscle cells are “scored” by low expression of *Nkx2-5* (LogTPM < 1, null expression) and high expression of smooth muscle cell genes (*Tagln, Cnn1, Acta2, Cald1, Mylk, Hexim1* and *Smtnl2* moderate to high (LogTPM >2) for at least 5 of these 7 genes). χ^2^ test: *p* < 2.37e-6. (**b**) Schematic illustration of forced expression of Nkx2-5 in Isl1^+^ cells and scRNA-seq. (**c**) Predicted diffusion pseudotime of Isl1^+^/Nkx2-5OE cells projected on t-SNE plots of Nkx2-5^+^ and Isl1^+^ (**d**) CPC.

### The landscape of chromatin accessibility in cardiac progenitor cells

To assess changes in genome-wide chromatin accessibility during early heart development, we performed ATAC-seq on FACS-sorted Nkx2-5^+^ and Isl1^+^ cardiac progenitor cells sampled at E7.5, E8.5 and E9.5 (Fig. 1b). For each condition and time-point, at least two biological replicates were used (Supplementary Table 4) (Supplementary Fig. 10a). In addition, we obtained transcriptional profiles of biological replicates by bulk RNA-seq at each corresponding developmental stage using the SMART-seq2 method [45] (Supplementary Fig. 10b). We detected a total of 5,866 differential chromatin accessibility peaks (log2[fold changes]>2 or <-2, FDR <0.05) in Nkx2-5^+^ cells and Isl1^+^ cells at different developmental stages using edgeR [46] for sequential pairwise comparisons of the Nkx2-5^+^ and Isl1^+^ samples. Interestingly, many differentially accessible regions in in Nkx2-5^+^ cells represented chromatin-opening events, although several chromatin closure events were also observed. In contrast, chromatin-closing events represented the vast majority of chromatin accessibility changes in Isl1^+^ cells (Fig. 5a) implying distinct chromatin states in either cell population. We observed robust closing chromatin peaks from E8.5 to E9.5 in the Nkx2-5 and the Isl1 lineage, suggesting that cell-fate restriction is associated with global loss of chromatin accessibility (Fig. 5a).

**Figure 5.**
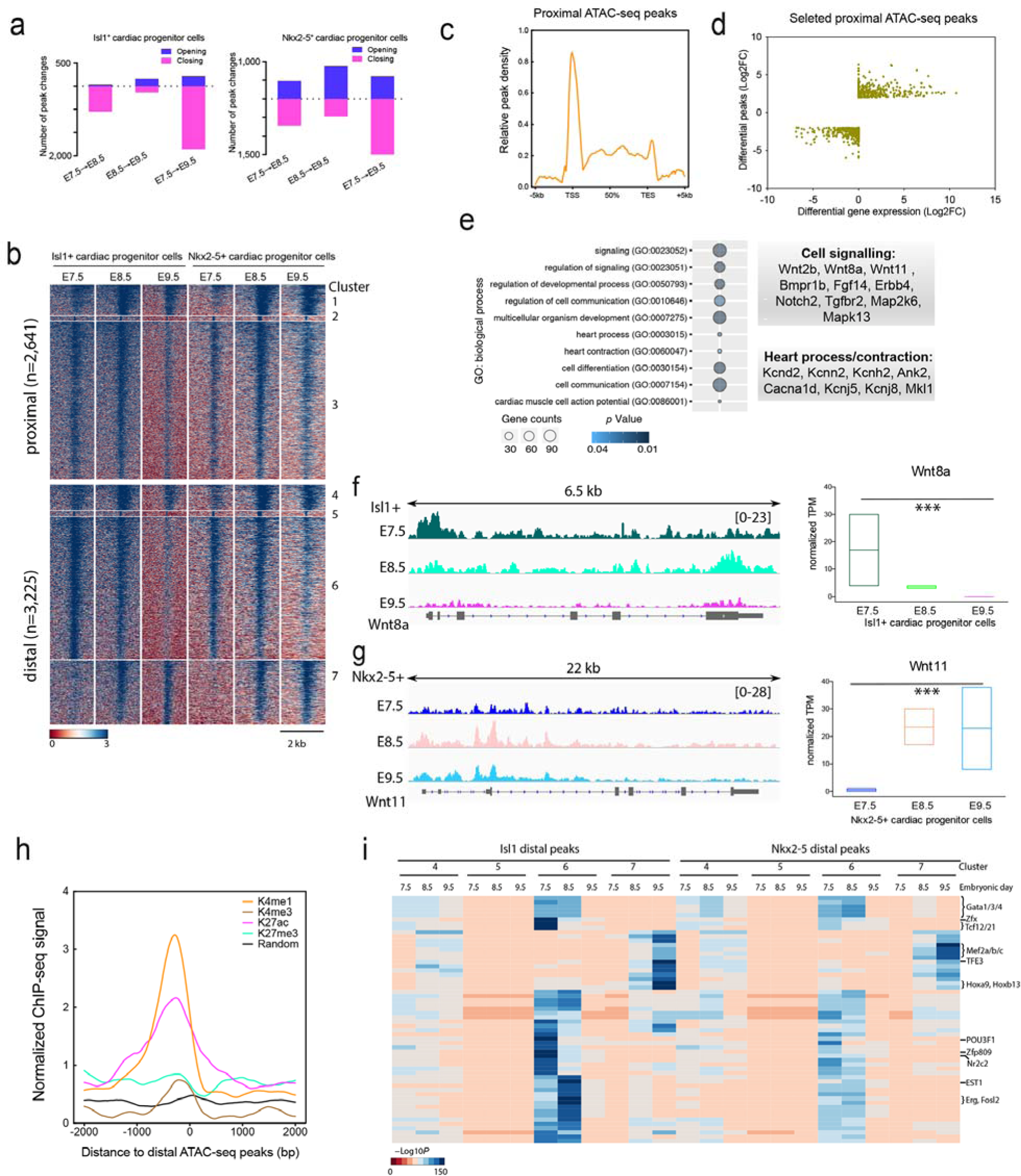
ATAC-seq based analysis of accessible chromatin and gene expression of cardiac progenitor cells. (**a**) Number of differential chromatin accessibility peaks in different transition states (log2(FC)>2, false discovery rate [FDR]<0.05). (**b**) Genome-wide distribution of differential open chromatin peaks grouped by distance to the promoters and by *K*-means. Each row represents one of 5,866 selected peaks in sequential comparisons (log2[FC]>2, FDR<0.05). (**c**) Metagene plot of proximal ATAC-seq peaks. (**d**) Plot showing accessible chromatin levels and gene expression levels of 805 selected differential-peak/DE-gene pairs. (**e**) Bubble chart showing enrichment of gene ontology terms of selected genes. (**f**) ATAC-seq profiles across Wnt8a and Wnt11 loci in Isl1^+^ and Nkx2-5^+^ CPC, respectively. (**g**) Boxplots showing gene expression of Wnt8a and Wnt11 in Isl1^+^ and Nkx2-5^+^ CPC. (**h**) Distribution of ChIP-seq reads for different histone modifications across distal ATAC-seq peaks in E10.5 embryonic hearts. (**i**) Heatmap showing enrichment of transcription factor motifs in each accessible chromatin cluster of Isl1^+^ and Nkx2-5^+^ CPC between E7.5 to E9.5.

*K*-means clustering of the genome-wide distribution of differential peaks revealed seven chromatin accessibility clusters for the three investigated developmental stages at locations proximal and distal to transcriptional start sites (TSS) (Fig. 5b). Cluster 1 and 7 showed increased chromatin accessibility from E7.5 to E8.5 whereas clusters 2-6 represented closing chromatin, probably associated with differentiation of CPC (Fig. 5b, Supplementary Fig. 11a). Between E8.5 and E9.5, we observed both a substantial loss of chromatin accessibility (cluster 6) and a gain of chromatin accessibility (cluster 7) (Fig. 5b, Supplementary Fig. 11a), which reflects the dynamic changes in chromatin accessibility associated with cell fate transition and terminal differentiation.

Next, we investigated whether changes in chromatin accessibility correlated with differential gene expression and which genes were subject to regulation. We detected 805 genes across all conditions for which differential appearance of opening peaks correlated with differential gene expression (Fig. 5c). Gene ontology (GO) analysis of this set of genes revealed enrichment for the cell signaling (Fig. 5d). A minor group of genes was associated with heart-specific term such as heart process (Fig. 5d). The genes involved in cell signaling comprised the *Wnt, BMP, FGF, Notch, TGF-*β and *Ras-MAPK* families suggesting that the dynamic changes in chromatin accessibility renders such cells responsive to environmental stimuli during development. For example, canonical Wnts like Wnt8a maintains proliferation of CPC proliferation while non-canonical Wnts (e.g. Wnt11) promote differentiation [47, 48]. Correspondingly, the ATAC-seq analysis demonstrated that the promoter of Wnt8a is closing from E7.5 to E9.5 during Isl1^+^ CPC differentiation (Fig. 5f) while the promoter and gene body of *Wnt11* are more accessible from E7.5 to E9.5 in Nkx2-5^+^ CPC (Fig. 5g).

ATAC-seq provides an excellent tool to identify transcription factor (TF) motifs that become accessible due to nucleosome eviction and/or chromatin remodeling [27]. Transcription factor binding often occurs at enhancers, which are marked by the histone modifications H3K4me1 and H3K27ac [49]. Concordantly, we noted that H3K4me1 and H3K27ac but not the repressive H3K27me3 were enriched at distal ATAC-seq peaks while random ATAC-seq peaks were devoid of H3K4me1 and H3K27ac (Fig. 5h). After exclusion of transcription factors that are not expressed in corresponding development stages according to our RNA-seq results, we scanned accessible peaks in chromatin for clusters 4-7 (distal peaks) at E7.5-E9.5, using the motifs from a set of 364 transcription factors and the motif analysis package HOMER [50]. Gata family TF motifs were enriched in cluster 6 both in Isl1^+^ and in Nkx2-5^+^ cells at E7.5 and 8.5 but not in clusters 4/5/7 and not at E9.5, suggesting that Gata TFs play roles in cell fate transition but not differentiation (Fig. 5i). Mef2a/b/c motifs were enriched at E9.5 in cluster 7 in both Isl1^+^ and in Nkx2-5^+^ cells, which corresponds to the known role of Mef TFs in cardiomyocyte differentiation [51]. Notably, we observed abrupt closure of sites containing motifs for several TFs including Zfx, Tcf12/21, Zfp809, POU3F1 and Nr2c2 from E7.5 to E8.5 in Isl1^+^ cells (cluster 6) indicating that binding of such TFs has to be abrogated quickly once CPC fate is established and before CPC specification occurs. Consistent with the loss of chromatin accessibility in cluster 6 from E8.5 to E9.5, most TFs found at E8.5 (cell transition state identified by our scRNA-seq as above) were completely absent at E9.5 in both Isl1^+^ and in Nkx2-5^+^ cells (Fig. 5i). Some TF motif patterns were conserved between Isl1^+^ and Nkx2-5^+^ cells, such as the pattern for Gata1/3/4, but the majority was different, suggesting that the same TF binds in a cell-type specific manner at different developmental enhancers to direct the fate of CPC. Taken together our newly established open chromatin atlas in combination with RNA-seq identified a set of transcription factors that seems to orchestrate early heart development through distinct developmental enhancers, which are differentially active in either Isl1^+^ or Nkx2-5^+^ cells.

### Isl1 and Nkx2-5 shape chromatin accessibility during cardiogenesis

To analyze how the lack of Isl1 expression affects chromatin accessibility, we performed ATAC-seq of Isl1 mutant CPC at E9.5. Inactivation of the *Isl1* gene resulted in comparable numbers of opening and closing peaks (386 opening and 345 closing) (Fig. 6a). Since our scRNA-seq analysis at E9.5 indicated arrest of Isl1 mutant CPC development, essentially converting the transcriptional profile of E9.5 into an E8.5 state (Fig. 3), we compared the landscape of chromatin accessibility of both populations. Surprisingly, we found that E8.5 Isl1^+^ cells and E9.5 Isl1 mutant cells show significantly different open chromatin signatures (432 opening and 728 closing), although they share similar positions at the developmental pseudotime trajectory (Fig. 6a), which indicates that Isl1-dependent changes in chromatin accessibility occur ahead of transcriptional divergence.

**Figure 6.**
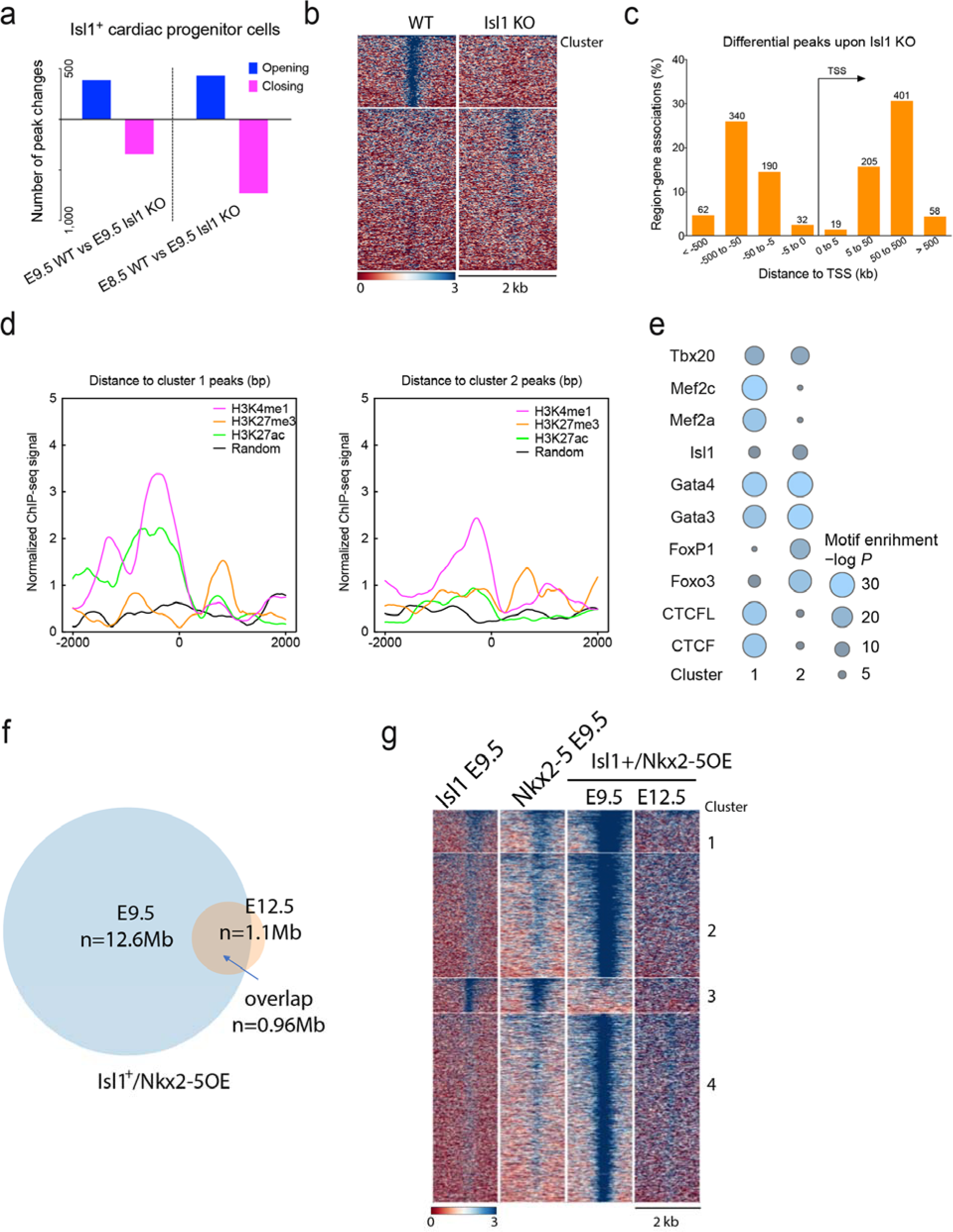
Dynamic changes in enhancer accessibility of Isl1 KO and Isl1^+^/Nkx2-5OE CPC. (**a**) Number of differential chromatin accessibility peaks in Isl knockout cells compared to wild type Isl1^+^ cells at E8.5 and E9.5 (log2[FC]>2, FDR<0.05). (**b**) Heatmap showing genome-wide distribution of differential open chromatin peaks grouped by *K*-means in Isl1 knockout and Isl1 control CPC. Each row represents one differential peak in pairwise comparison (log2[FC]>2, FDR<0.05). (**c**) Bar plot showing the distance of differential peaks to the nearest promoters. (**d**) Distribution of ChIP-seq reads for histone modifications across cluster 1 (left) and 2 (right) ATAC-seq peaks in E10.5 embryonic hearts. (**e**) Bubble chart showing the enrichment of transcription factor motifs in each cluster of differential peaks. The color and area of circles represent the *p-*value. (**f**) Venn diagram showing overlap of genome coverage of open chromatin peaks between E9.5 and E12.5 Isl1^+^/Nkx2-5OE cell. (**g**) Heatmap showing genome-wide distribution of differential open chromatin peaks grouped by *K*-means in Isl1^+^/Nkx2-5OE cells at E9.5 and E12.5 in comparison with wildtype Nkx2-5^+^ and Isl1^+^ cells.

Loss of Isl1 led to more robust opening than closing of chromatin regions as identified by *k*-means clustering (cluster 1 peaks depend on Isl1 for opening as they are closed upon Isl1 knock-out; cluster 2 peaks depend on Isl1 for closing) (Fig. 6b, Supplementary Fig. 11c), which suggests that Isl1 is required to leave the attractor state characterized by an open chromatin organization. Annotation of opening and closing peaks by GREAT analysis [52] showed that 96.2% of the peaks (1,256) located in distal regions (> ±5 kb to the TSS sites) presumably representing enhancer regions affecting cardiac progenitor cell decisions (Fig. 6c). Both cluster 1 and 2 peaks showed enrichment of H3K27ac and H3K4me1 but not of H3K27me3 supporting this conclusion (Fig. 6d). Investigation of transcription factor motifs enriched in either opening or closing peaks using the motif analysis package HOMER [50] revealed that binding sites for GATA family factors and Tbx20 are enriched in both opening and closing peaks, suggesting an Isl1-independent mode of action. In contrast, binding sites for Mef2-family members were only enriched in Isl1-dependent opening peaks while binding sites for Forkhead box family TFs were enriched in Isl1-dependent closing peaks (Fig. 6e). We concluded that Isl1 acts together with Mef2 factors but prevents binding of Forkhead factors to guide cardiac progenitor cell fate decision. The enrichment of CTCF and CTCFL in open peaks implied that Isl1 alters the topology of the chromatin to achieve chromatin opening. Surprisingly, Isl1 binding motifs were not significantly enriched in either opening or closing peaks suggesting that Isl1 changes chromatin accessibility not directly but by secondary effects, e.g. CTCF/CTCFL mediated-chromatin reorganization. In contrast to the inactivation of Isl1, which led to more robust but moderate number of opening peaks, forced expression of Nkx2-5 clearly increased chromatin accessibility at multiple loci, compared to either Nkx2-5^+^ or Isl1^+^ cardiac progenitor cells, and few closing peaks (opening peaks: 556 for Nkx2-5^+^ and 1,526 for Isl1^+^; closing peaks: 73 for Nkx2-5^+^ and 86 for Isl1^+^) (Supplementary Fig. 11c). Interestingly, forced expression of Nkx2-5 resulted in a dramatic increase of accessible chromatin sites at E9.5 compared to E12.5 and Nkx2-5^+^ or Isl1^+^ CPC at E9.5 (Fig. 6f, g). However, we noted that chromatin opening occurred only transiently but was not sustained upon CPC differentiation (Fig. 6f). We hypothesized that chromatin opening evoked by Nkx2-5 overexpression was overcome by cellular events set in motion by Nkx2-5 during differentiation. Similar to Isl1, most effects of forced Nkx2-5 expression were confined to distal regions (96.7% of 3,374 peaks) probably representing enhancers (Supplementary Fig.11e). Motif analysis of these peaks revealed that most peaks contained Sox family factor binding sites. Surprisingly, the Nkx2-5 motif was not significantly enriched suggesting that the altered chromatin accessibility caused by forced Nkx2-5 expression is mediated via secondary effects (Supplementary Fig. 11f).

## DISCUSSION

Our scRNA-seq analysis provides a rich data source for the discovery of genes that might play a role in heart development. For example, we found that posterior *Hox* genes are temporarily expressed in early stage Isl1^+^ cells. *Hox* genes are well known to establish the anterior–posterior axis during embryogenesis [53] and anterior *Hox* genes (*Hoxa1*, *Hoxb1*, and *Hoxa3*) are involved in cardiac development [54–56] but expression of posterior *Hox* genes had not been detected in cardiac progenitor cells so far. We speculate that posterior Hox family transcription factors might contribute to patterning of the heart, which is supported by the presence of cardiac defects in *Hox A/B* cluster compound mutants [57], However, to unravel specific functions of posterior *Hox* genes for cardiac morphogenesis it might be required to concomitantly inactivate several paralogous posterior *Hox* genes, thereby avoiding potential compensatory effects, which are probably responsible for the absence of cardiac phenotypes in individual posterior *Hox* genes mutants (e.g. Hoxb6) [58].

Lineage tracing and clonal analyses have demonstrated that Nkx2-5^+^ cardiac progenitor cells contribute to cardiac endothelium and smooth muscle cells [43]. However, we did not detect Nkx2-5 expression outside of the cardiomyocyte lineage during differentiation, which seems to be in conflict with lineage tracing studies but is consistent with prior studies allocating Nkx2-5 primarily to cardiomyocytes [21]. It is important to remember that tracing or actual reporter-based gene activity approaches address different questions. In our study, we exclusively focused on Nkx2-5 expressing cells but not on their derivatives, which excludes Nkx2-5-derived cells that have terminated Nkx2-5 expression. We believe that the loss of a bipotent fate of Isl1^+^ cells and the acquisition of a unipotent cardiomyocyte fate after forced expression of Nkx2-5 clearly argue for a decisive role of Nkx2-5 in cardiomyogenic differentiation. In agreement with this hypothesis, we observed a concomitant increase of cardiomyocyte genes such as Myl7 [42, 59] together with Nkx2-5 expression at early development stages (Fig. 1f, 2e) as well as rapid differentiation of progenitor cells to smooth muscle cells once Nkx2-5 expression declined.

So far dynamic changes in the genome-wide chromatin landscape have not been systematically investigated during early heart development, although chromatin remodeling has been linked to heart development and the BAF chromatin-remodeling complex was identified as a crucial factor [59, 60]. Our ATAC-seq data covers ~1.5% of the mouse genome (data not shown) and theoretically covers most of the active *cis-*regulatory elements, thereby enabling precise views on open chromatin sites associated with distinct cell fates and identities during early cardiac development. We found that inactivation of Isl1 dramatically changed the accessibility of CTCF and CTCFL binding sites, which most likely will alter topology associated domains (TAD) in the chromatin [61].

ATAC-seq peaks cover numerous different genomic elements such as enhancers, insulators and promoter, which make a clear association of open chromatin states with gene expression difficult. We therefore only correlated proximal peaks around transcriptional start sites with gene expression, which resulted in the identification of two different GO categories, i.e. (i) cell signaling and (ii) heart process. Both GO categories correlated well with the different developmental stages and reflected the respective stages of CPC differentiation.

Analysis of distal peaks, which we deliberately did not link to close or more distant active genes, enabled us to analyze putative dynamic changes in enhancers that act as integration hubs for transcription factors [49]. Motif enrichment analysis of distal peaks, which comprised 55% of all differentially accessible peaks, allowed identification of binding sites for several TFs such as POU3F1 and Nr2c2, which so far have no clearly assigned roles during heart development. Furthermore, loss/gain of function studies suggested that the two cardiac transcription factors Nkx2-5 and Isl1 execute their functions by regulating chromatin accessibility for similar sets of transcription factors but in a cell specific manner at different developmental enhancers.

The power of our combined approach to identify potential new regulatory circuits might be illustrated using the example of posterior *Hox* genes. The scRNA-seq analysis indicated that *Hox* family genes are expressed at early development stages but corresponding binding motifs in open chromatin regions were only identified at later stages. Since development and differentiation of CPC occurs within a narrow window (E7.5-E9.5, 48 hours), it seems likely that the Hox proteins at this stage are subject to posttranslational regulation and show an extended half-life to execute functions during CPC differentiation. The integration of different regulatory layers into a comprehensive model that explains different developmental decisions during heart development will be a major challenge for the future. The complex network of stage-specific transcription factor enhancer interactions and the single cell transcription profiles revealed in our study provides part of the essential groundwork to move in this direction.

## METHODS

### Mouse work and sampling of single cells

All animal experiments were carried out following regulations of the Committee for Animal Rights Protection of the State of Hessen (Regierungspraesidium Darmstadt, Darmstadt, Germany). The transgenic mouse lines used in this study have been described previously [9, 10]. C57BL/6 mouse embryos were dissected at E7.5, E8.5, E9.5 or E12.5. Embryonic hearts were isolated under the dissection microscope and digested into single cells suspensions with 0.25% trypsin-EDTA. After washing with PBS, cells stained were with DAPI to check for viability, and sorted using the GFP channel of the BD FACSAria II instrument. To obtain *Isl1^−/−^* or Isl1^+^/Nkx2-5OE cells, *Isl1^nGFP/+^* mice or *Isl1-Cre* and *Rosa26*^Nxk2-5-IRES-GFP^ mice were mated and embryos were recovered at indicated time points. Genotyping was achieved by PCR using non-heart tissue of the same embryos as described [10].

### Single-cell RNA sequencing library preparation

Single-cell capture, lysis, reverse transcription, and pre-amplification was done using C1 chips (#100-5763, 10-17 μm) in the C1 single-cell Auto Prep System (Fluidigm) or the ICELL8™ Single-Cell System (Wafergen) following the manufacturers protocols. Libraries were sequenced using an Illumina NextSeq 500 system.

### Single-cell RNA-seq data analysis

#### 1. Raw data processing

Low quality bases were trimmed off the raw sequencing reads using Reaper with a minimum median quality of 53 in a window of 20 bases, omitting the first 50 bases of the read. Additionally, the ‐dust-suffix 20/AT option was used to trim remaining polyA or polyT stretches at the end of reads as well as stretches of B (a special Illumina Quality Score indicating non-thrustworthy bases) with the –bcq-late option. The STAR alignment tool was used with default parameters to map trimmed reads to the mouse genome (version mm10) and transcriptome (–quantMode TranscriptomeSAM, together with the Gencode annotation in version vM10). Mapping quality and statistics was assessed using Qualimap in rnaseq mode, setting the protocol to strand-specific forward and using the same Gencode annotation. The Qualimap output was used later for single-cell filtering (see below). RSEM was used with gene annotations from Gencode vM10 as well as a single-cell prior to assign reads to genes and extract gene-centered counts.

#### 2. Cell quality and filtering

A SingleCellExpression-Set object (SCESet, R package scater) was created in R from all available metadata, cell quality data, gene annotations and the gene-centered count table. For each platform (Fluidigm C1 (C1) or Wafergen (WG)), an initial cell-quality map was generated with t-SNE (R package Rtsne) by grouping cells with similar quality metrics together (Supplementary Figure 1). The (per-cell) quality metrics used as input were: number of features (genes) detected with at least 10 counts, the percentage of gene dropouts, the number of alignments, the number of alignments to exons, introns and intergenic regions, the number of secondary alignments, the expression of Rplp0 (also known as 36B4) as housekeeping gene, the percentage of read counts to mitochondrial genes, as well as the percentage of genes detected.

To define cells as low quality, we formulated and evaluated five criteria for each cell: The percentage of counts to mitochondrial genes is 1.5 median-absolute-deviations (MADs) above the median, the number of detected features is 2 MADs above or below the median, the percentage of gene dropouts is 2 MADs above the median, the Rplp0 expression is 2 MADs below the median and the percentage of genes is 1.5 MADs above or below the median. Cells failing more than one criterion were considered low quality and excluded from further analysis. See Supplementary Fig. 1 for a graphical representation of cell filtering.

#### 3. Gene filtering and expression normalization

Similar to cell filtering, we defined two criteria for gene filtering: (1) gene expression across all cells of a lineage (excluding cells from knockout and overexpression experiments) exceeds 2000 counts and (2) at least 10 cells from a lineage show gene expression above 10 counts. A gene was filtered if it failed at least one criterion in both lineages. After filtering, count data of 12053 genes across 498 cells remained for further downstream analysis. Remaining count data were normalized by separately applying the sum factor method, as implemented in the R package scater, to cells from the two lineages.

#### 4. Preprocessing of single-cell data of Nkx2-5 knockout hearts

We combined count tables obtained from wildtype cardiac single cells across time points E8.5, E9.5 and E10.5, as well as from Nkx2-5 knockout cardiac cells [20] from E9.5 into a single SCESet object and filtered out cells that were identified as low-quality. After filtering, count data from 11781 genes across 2358 cells were used to cluster cells using the quickCluster command from the R package scran. Sum factor normalization was applied with deconvolution of size factors within obtained clusters.

#### 5. Definition of heterogeneous genes

Sum factor normalized counts were used to define heterogeneous genes within lineages as well as at individual time points. Specifically, we calculated the coefficient of variation as well as the dropout-rate per gene and investigated their relationship to the mean expression of that gene. We next binned both (ordered) statistics into windows of size 200 and scaled values (z-score transformation) within windows. Genes for which one of the scaled statistics exceeded a 99-percentile within its window where called heterogeneous.

#### 6. Clustering

We scaled normalized expression values of heterogeneous genes and used them as input to dimension reduction by self-organizing maps (SOMs) for each lineage. Briefly, SOMs or Kohonen Networks were treated as special cases of neuronal networks, where no target vector containing class labels is necessary for training. Instead, a map is initialized randomly for each cell, consisting of fewer map tiles than input genes, effectively representing meta genes. During training, genes are subsequently placed onto map tiles with the most similar meta gene representation. Importantly, a gene ends up on the same map tile of all cell maps, therefore creating a lower dimensional representation of the cell’s transcriptome using meta genes. After 2000 training epochs, cell maps were further projected into two dimensions by t-SNE (perplexity value of 15, 2000 epochs of convergence) and clustered with HDBSCAN using a minimum cluster size of 7 and min_samples 9 (Fig. 1c, d and Supplementary Fig. 2a, b).

#### 7. Differential expression and lineage dynamics

Differentially expressed genes between cell clusters were assessed using MAST on sum factor normalized counts (log2 scale). The MAST framework models gene expression in a two-component generalized linear model, one component for the discrete expression rate of each gene across cells and the other component for the continuous expression level, given the gene is expressed. Additionally, we used a gene ranking approach (SC3) to define marker genes specific for each cluster (Supplementary Table 1, 2). To define lineage dynamics, we used all protein coding genes that were marker genes for a cluster (AUROC > 0.8, FDR < 0.01) and differentially expressed in any cluster (lower bound of LFC > 2 or higher bound of LFC < −2, FDR < 0.01) as input to destiny (Fig. 2a, b).

#### 8. Cell transition states and transcriptome noise

For the critical transition index (*I_C_(c)*), we computed the absolute marker gene-to-gene and cell-to-cell correlations for each cluster and calculated the ratio of their means (Supplementary Fig. 6a, b). To reduce influence from differing cell numbers in clusters, we applied a bootstrapping procedure, randomly selecting 30 (20) cells from a given Nkx2-5 lineage cluster or Isl1 lineage cluster repeating the procedure 1000 times. Pairwise cell-to-cell distances were calculated as described by Mohammed and colleagues[41].

#### 9. Gene correlation analysis

To define gene networks that play a role in lineage development, we assumed that genes expression will either increase or decrease with lineage progression. Therefore, we calculated the (global) Spearman’s Rank correlation of the expression of each gene to the diffusion pseudotime from destiny. Since a gene might exhibit its expression dynamics only within discrete states (clusters), we also calculated the (local) Spearman’s Rank correlation of gene expression to pseudotime within clusters. We defined a gene as correlated gene, if it shows a global correlation of at least 0.7 or a local correlation of at least 0.5 (Supplementary Table 3). Lineage-specific correlated genes were used to identify gene networks. Genes within the same sub-network show a high correlation (measured as Pearson’s Correlation), but a lower correlation between sub-networks (Supplementary Fig. 7a, b). To identify the dynamics of correlated genes, expression was smoothed along pseudotime by calculating the mean expression in windows of 11 consecutive cells (Supplementary Fig. 7 c, d).

#### 10. Correction of batch effects

To join data sets from two different sequencing platforms, normalized expression values from heterogeneous genes were used as input into the mnnCorrect function from the R package scran. Briefly, mnnCorrect finds cells from different platforms that have mutually similar expression profiles. This is done by identification of pairs of cells that are mutual nearest neighbors, which can be interpreted as belonging to the same cell state. For each MNN pair, the method estimates a pair-specific correction vector. Those vectors are in turn averaged with nearby MNN pair vectors from the same hyperplane using a Gaussian-Kernel to obtain more stable cell-specific correction vectors. The procedure allows correction of cells that are not part of any MNN pair, e.g. data set specific cells that were sampled only on one platform. Corrected expression values were used for clustering and differential expression analysis analogous to steps 6 and 7 (Supplementary Fig. 5).

#### 11. Cell cycle scores of single cells

Cell cycle scores were calculated for each known cell cycle stage (G1/S, S, G2, G2/M, M/G1) using gene sets described by Whitfield et. al [62]. Specifically, a raw score was calculated as the average expression of genes in each set. To refine the score, we determined genes that correlated (rank correlation > 0.4) well with the raw score and calculated the cell cycle score using those genes. Cell cycle scores were z-score transformed (scaled) before plotting. A test of equal proportions was then conducted for cycling cells among Isl1^+^ and Isl1^-/-^ cells.

### ATAC-seq library preparation and sequencing

2, 000-20,000 of GFP^+^ cardiac progenitor cells were FCAS-purified and used for ATAC-seq. The ATAC-seq libraries were prepared as previously described [27]. 2×50 paired-end sequencing was performed on Illumina NextSeq500 to achieve on average of 36.4±12.7 million reads per sample.

### ATAC-seq data analysis

#### 1. Raw data processing

Raw ATAC-seq paired-end reads were trimmed and filtered for quality, and then aligned to the mouse genome GRCm38 (mm10) using STAR [63] with the following parameter: –outFilterMismatchNoverLmax 0.2 –outFilterScoreMinOverLread 0 –outFilterMatchNminOverLread 0 –outFilterMatchNmin 20 –alignIntronMin 2 –alignIntronMax 1 –outFilterMultimapNmax 1 –alignMatesGapMax 2000 –alignEndsProtrude 10 ConcordantPair. Reads that did not map, non-uniquely mapped, mapped to repetitive regions and chromosome M, and PCR duplicates were removed.

#### 2. Normalization, peak calling and open chromatin atlas generation

For downstream analysis, the read counts were normalized to 1× depth (reads per genome coverage, RPGC) using the “bamCoverage” function of deepTools2 [64]. Peak calling was performed using “callpeak” function of MACS2 [65] with the following parameters: –nomodel –shift −100 –extsize 200 -q 0.05. Peaks in each sample were merged as union peaks for calculation of peak counts and quality control purposes. The normalized number of reads mapped to each peak of the union peaks in each sample was quantified using bigWigAverageOverBed (https://github.com/ENCODE-DCC/kentUtils). Peak counts of all samples were then merged to obtain a data matrix and normalized with edgeR [46]. To remove the non-reproducible replicates, we calculated Pearson correlations using log2 normalized counts and removed pairs, in which the Pearson correlation was below 0.8 resulting in at least two replicates for each developmental stage. Differential accessible peaks were pairwise-compared sequentially across each developmental stage and combined as a genome-wide atlas of the accessible chromatin landscape in CPC.

#### 3. Genome-wide distribution of differential peaks

The normalized read counts for each developmental stage across replicates were merged, binned around all differential peak summits in 50 bp bins spanning ±1 kb region, clustered by *k*-means algorithm, and visualized by creating heat maps using deepTools2 [64].

#### 4. Assignment of proximal and distal ATAC-seq peaks

The proximal and distal peaks are defined by the distance of differential ATAC-seq peaks towards annotated promoters (Gencode annotation): at least 2.5 kb away from promoters were selected as distal peaks, and the others were assigned as proximal peaks.

### Bulk RNA-seq

5,000-20,000 of cardiac progenitor cells were sampled using the same protocol as described above for scRNA-seq. Bulk RNA-seq libraries were prepared using Smart-seq2 according to the manufacturer’s protocol (#634889, Clontech), and sequenced using the Illumina NextSeq500 instrument. Raw reads were processed using the same method as for scRNA-seq. Quantification and identification of differentially expressed genes were carried out using DEseq2 [66].

### Statistics

Divergence from the median absolute deviation (MAD) of several quality metrics was used to identify outliers in single-cell data. Differential expression analysis between clusters of single cells was evaluated using a two-component Hurdle model as implemented in the R package MAST. Likelihood ratio (LR) tests for combined testing of differences in proportion of expression and average expression were applied and obtained p-values were corrected for multiple testing using false discovery rates (FDR). P-values for marker genes were calculated using a signed Wilcoxon rank test and corrected for multiple testing using FDR. Differences of proportions (e.g. proportions of different cell types) were assessed with a test of equal or given proportions, as implemented in the R function prop.test. Differential accessibility was calculated from ATAC-seq data by evaluating a General Linear Model (GLM) and tested using LR tests as implemented in the bioconductor package edgeR. Obtained p-values were also corrected for multiple testing using FDR. No statistical methods were used to predetermine sample size. The experiments were not randomized. The investigators were not blinded to allocation during experiments and outcome assessment.

### Code availability

The R scripts used for data analysis and simulations are freely available on request.

### Data availability

All raw and processed data are freely available from the ENA repository (https://www.ebi.ac.uk/ena) under accession number PRJEB23303.

## AUTHOR CONTRIBUTIONS

G.J. and T.B. designed and conceived the project. J.P. and G.J. analyzed the scRNA-seq data. G.J. analyzed the ATAC-seq data. G.J. isolated and dissected embryos. G.J., S.G. and M.Y. performed scRNA-seq and ATAC-seq. X.Y. provided transgenic mouse lines. S.G., C.K., M.L. and Y.Z. contributed to data processing, discussions and advice. G.J., J.P. and T.B. wrote the manuscript.

## Acknowledgements

We thank Ann Atzberger for help with cell sorting and Hui Qi for assisting with isolation and dissection of mouse embryos. This work was supported by the Excellence Initiative ‘‘Cardiopulmonary System’’ (ECCPS), the DFG collaborative research center SFB1213 (TP A02 and B02) the Foundation Leducq (3CVD01), the German Center for Cardiovascular Research and the European Research Area Network on Cardiovascular Diseases project CLARIFY.

